# Optimal Conservation Prioritization for Waterfowl and Other Birds in Eastern Canada

**DOI:** 10.64898/2026.10.30.763193

**Authors:** Daniel Dylewsky, Matthew E. Dyson, James E. Paterson, Richard Schuster, Joseph R. Bennett

**Affiliations:** Department of Biology, Carleton University, 1125 Colonel By Drive, Ottawa, K1S 5B6, ON, Canada.; Institute for Wetland and Waterfowl Research, Ducks Unlimited Canada, 1 Mallard Bay, Stonewall, R0C 2Z0, MB, Canada.; The Nature Conservancy of Canada, 365 Bloor Street East, Suite 1501, Toronto, M4W 3L4, ON, Canada.

## Abstract

Conserving and restoring high-quality habitat is widely recognized as one of the most effective means of supporting migratory and wetland bird populations. To advance this objective at a regional scale, the Eastern Habitat Joint Venture (EHJV) coordinates conservation planning for 22 focal bird species across Canada’s eastern provinces. Optimization approaches to support conservation planning are increasingly common, and can be especially valuable in large areas where resources are limited. We present an optimized geospatial prioritization scheme designed to identify regions of primary importance to species coverage objectives within the EHJV. We apply linear programming methods to species abundance data products to maximize population coverage using an adaptable variable-target approach, whose outputs generalize readily to delivery projects of any budget or coverage goal. Our base results highlight areas of high conservation value for priority waterfowl and non-waterfowl species, and we present additional examples of how region-wide solutions can be subdivided based on jurisdictional boundaries, land ownership, or existing protected status. We also address methodological decision points, with attention to how available species abundance data sources were evaluated and what considerations were involved in formulating the optimization computations. We emphasize possibilities for future work to incorporate additional conservation values, such as carbon storage and flood mitigation, and costs, such as land value, and feasibility.

## 1 Introduction

Georeferenced prioritization schemes developed using quantitative optimization methodologies play an increasingly central role in conservation planning for habitats and wildlife populations. The proliferation of spatially explicit data describing species distributions, habitat conditions, threats, and conservation opportunities has enabled conservation organizations to make decisions based on transparent and reproducible objectives. This broader discipline of systematic conservation planning (SCP) seeks to identify spatial configurations of conservation actions that maximize ecological benefit while minimizing fiscal or social costs (Margules and Pressey 2000; McIntosh et al. 2017).

Decision support tools built on an SCP framework have played an increasingly important role in questions of resource allocation for conservation planning due to their ability to offer transparent and reproducible geospatial prioritizations with objectives reflecting species’ habitats (e.g., Ball et al. (2009); Moilanen et al. (2022); Hanson et al. (2025)), threat-based targets (e.g., Carwardine et al. (2019)), or species persistence (e.g., Joseph et al. (2009); Bennett et al. (2014)). Many conservation prioritization efforts rely on linear value modeling approaches that rank locations according to weighted combinations of species and habitat attributes (e.g., Krainyk et al. (2019); Couvillon et al. (2026)). More recently, advances in SCP have shifted emphasis toward optimization-based approaches that explicitly account for complementarity among sites and identify conservation portfolios that collectively achieve management objectives more efficiently (Kukkala and Moilanen 2012; McIntosh et al. 2017). These advances have led to widespread adoption of planning tools such as Marxan (Ball et al. 2009), Zonation (Moilanen 2007), and mixed-integer linear programming (MILP; Beyer et al. (2016); Schuster et al. (2020)). Despite these advances, conservation planning often remains constrained by uncertainty in ecological data, changing conservation priorities, and the practical reality that budgets, partnerships, and implementation opportunities vary considerably across organizations and jurisdictions. Consequently, there remains a need for prioritization frameworks that maintain the rigor of optimization-based approaches while producing flexible decision-support products that can be adapted to a wide range of planning contexts.

This paper presents an SCP approach for bird species within Canada’s Eastern Habitat Joint Venture (EHJV). The EHJV is one of the largest habitat Joint Ventures established under the North American Waterfowl Management Plan (NAWMP; North American Waterfowl Management Plan (2024)). The EHJV spans six eastern Canadian provinces and supports conservation planning across more than 3 million km^2^. Conservation Priority Areas currently used by the EHJV were originally developed using information and analytical approaches that predate major advances in species distribution modeling, remote sensing, optimization methods, and the understanding of climate change and other emerging conservation challenges. In response, the 2021-2030 EHJV Implementation Plan identifies the development of an updated landscape prioritization framework as a key objective (Eastern Habitat Joint Venture 2024). Decision support tools have become a hallmark of NAWMP implementation and are increasingly used to translate continental conservation objectives into spatially explicit priorities for habitat conservation. Collectively, these efforts reflect a progression from structured, multi-objective spatial value models (Krainyk et al. 2019) toward increasingly flexible and scalable approaches for conservation planning. These frameworks include linking local conservation delivery to regional and continental objectives (Palumbo et al. 2021), evaluations of alternative approaches for incorporating species distributions into spatial prioritizations based on expert opinion (Winiarski et al. 2024), regional planning tools that allow priorities to be tailored across scales and stakeholder objectives (Couvillon et al. 2026), and waterfowl distribution and productivity products developed to support conservation planning in the Prairie Pothole Region (Devries et al. 2023; Prairie Habitat Joint Venture 2021).

The decision support product developed in this paper is designed to update EHJV priority areas by incorporating new data sources, expanded species considerations, and improved optimization methodologies. In an effort to ensure that these results are useful to all conservation practitioners, regardless of the geographical scope or fiscal budget of their work, we have devised a methodology to produce maps which assign priority scores (from 0 to 1) across the full extent of the EHJV, divided into a grid of planning units made up of hexagons 10 km wide. These scores reflect the estimated conservation value of each grid cell, based on the relative local abundance of priority species. They are computed based on the results of optimization routines which solve for the most area-efficient spatial configurations that meet designated species coverage targets. We use a mixed-integer linear programming (MILP) approach capable of exactly solving the defined optimization problems. Such optimizations can also be carried out through simulated annealing methods or modified greedy algorithms (popularly implemented in the conservation planning packages Marxan (Ball et al. 2009) and Zonation (Moilanen et al. 2022), respectively), but modern exact integer linear programming solvers can offer more transparent and interpretable results without need for calibration (Schuster et al. 2020). Here, we integrate contemporary species distribution models for priority waterfowl and non-waterfowl species with a MILP approach to develop reproducible conservation priorities for the EHJV. To enhance their utility across a broad range of conservation decisions, optimization results are translated into gradated prioritization products rather than binary reserve designs, providing a flexible and adaptive decision-support framework that can readily incorporate new data, evolving conservation objectives, and future planning scenarios.

Our planning scheme is based on existing species distributions: we assume that conservation delivery will offer the greatest benefit in regions where priority species are abundant. The data layers representing these species distributions are drawn from published species abundance products, without attempting to improve upon (or synthesize) existing models (which falls outside the scope of this study). While the EHJV places primary conservation emphasis on waterfowl, its current Implementation Plan specifies 16 priority non-waterfowl species whose representation should be considered alongside the 6 priority waterfowl species. In order to accommodate this expanded scope, we produced prioritization maps for waterfowl and non-waterfowl priority species separately, with the intention that the reader can cross-reference the two recommendation schemes. The data processing and optimization methodology we present (and the accompanying code base) are modular and adaptable. The prioritization results we have produced are designed to serve current conservation priorities using the best available data, but priorities inevitably shift over time, and new datasets arise. Our approach is intended to generalize readily to evolving conservation needs, whether that means using updated abundance data products, alternate species inclusions, or a modified spatial domain. Our goal is not to provide a single, finalized prioritization product, but to demonstrate a workflow that can be used to continually iterate and improve on our results through a structured decision-making process with appropriate stakeholder engagement (as in Robinson et al. (2016), e.g.). For example, we demonstrate multiple nodes where additional expert information can be solicited and formalized into the process including species selection, species weighting, and seasonal selection.

## 2 Methods

### 2.1 Species Abundance Data Sources

The central data requirement of our analysis is spatial predictions of species abundance which cover the priority species and geographic extent of the EHJV. We considered a number of candidate data products and elected to use the 2023 relative abundance layers published by eBird Status and Trends (Fink et al. 2025), based on species distribution models trained on citizen science data. This decision was primarily motivated by species coverage: the eBird product was the only candidate data source that contained the 6 priority waterfowl species and 16 priority non-waterfowl species which are the focus of this analysis. No other source covered both waterfowl and non-waterfowl, and even within those categories they all lacked some priority species. Moreover, the eBird layers offer better spatial coverage of the EHJV area than the other candidate sources. eBird data for some species are absent in the more northern latitudes, but data scarcity in these regions is an issue common across all available data sources. The benefits of the eBird products have led to increasing adoption by joint venture partners across North America (Farwell et al. 2025). The alternative abundance data sources that we considered were those published by Frei et. al. (Frei et al. 2024) and Adde et. al. (Adde et al. 2020a). The Boreal Avian Modelling Centre’s Landbird Models also offer georeferenced avian abundance data within the study area (Solymos et al. 2025; Stralberg et al. 2025), but were not considered for this analysis because they do not include any of the waterfowl or waterbird species of central concern. We present a full breakdown of species and area coverage in the supporting materials (Table E1, Fig. E12).

Direct comparison of the three best candidate sources for waterfowl data reveals some discrepancies. While there are many coarse similarities across the data sources, each exhibits some features which are not well reproduced by the others. Without any ground truth against which to compare, it is difficult to directly assess the quality of each source. Aggregating data from multiple sources (e.g., by averaging) for the purpose of prioritization analysis is not a trivial task. Each source reports relative abundance on a different scale, and even if they were rescaled to share a common mean, their distributions would still be quite different from one another. For the prioritization problems with which we are concerned, the distribution of lower-valued grid cells is less important; the fact that the eBird layer sets many more values to zero than the other sources, for example, has little effect on prioritization schemes which target high-abundance regions. Significant disparities in the sources’ placement of high species abundance, however, should be cause for concern. Although we believe the level of agreement across available data sources is sufficient to justify their use as a foundation for data-driven conservation prioritization planning, the differences that they do exhibit highlight the need for continued improvement in the observation and modeling procedures used to estimate relative abundance.

### 2.2 Planning Units

Development of a spatial conservation prioritization scheme requires the division of the study area into a geographical grid of planning units. One possible approach would be to choose grid units which reflect how conservation action is delivered. In this spirit, we considered hydrological divisions (at the sub-watershed level) and ecological divisions (using existing ecodistrict classifications (Ecological Stratification Working Group 1996)). Both of these methods offer the advantage of corresponding to divisions which may already be used in determining jurisdictions of conservation delivery, as well as providing a hierarchy of planning unit size—in general, units are more densely packed in regions whose topography or ecology exhibits more heterogeneity.

Unfortunately, the use of these context-informed divisions has its drawbacks. Planning units of non-uniform size can offer greater planning resolution in regions of interest, but can also hinder interpretation of results (Pressey and Logan 1998; Nhan-cale and Smith 2011). Cells can be evaluated based on absolute conservation benefit valuation or on a per-unit-area basis, but either approach can lead to unwelcome consequences: smaller units are more likely to become outliers, while larger units may wash out localized variations within their perimeters. This results in a planning scheme whose distribution is influenced considerably by the choice of cell boundaries, which is not ideal.

Furthermore, many of the watershed and ecodistrict divisions have boundaries along coasts, which can lead to particular difficulty when analyzing abundance distributions of species which tend to have high concentrations along shorelines. Since rasterized abundance data is generally reported on a uniform square grid, aggregating it into zones whose boundaries are likely (by design) to transect regions of high abundance results in a high incidence of population centers being awkwardly split across planning units.

In order to mitigate these challenges, we have conducted our analysis on a regular grid. Although this does sacrifice the logistical naturalness of the ecodistrict and watershed schemes, it offers the benefit of uniform cells of equal area whose boundaries do not (except by chance) correspond to underlying features in the data. We have chosen to use a grid of regular hexagons 10 km across (i.e., with side length 5.77 km), which offers a higher spatial resolution than the other candidate schemes. This enables more fine-grained interpretation of results, while still allowing for subsequent coarse-graining: results on the hex grid could later be aggregated to the ecodistrict level, for example. The capacity for higher spatial granularity is particularly useful for species which form highly localized population centers, such as on shorelines or riverways. The grid scale could have been further reduced to provide even finer resolution—a logical lower bound is provided by the 3 km square grid on which the abundance data is reported. The chosen 10 km grid represents a compromise designed to leave greater room for local expertise at the site-specific project level. After removing cells that were all open water, our study area in the EHJV contained 37195 hexagonal cells (each of area 86.6 km^2^).

### 2.3 Formulating the Spatial Prioritization Problem

The goal of this study was to provide a data-driven geospatial prioritization of the conservation value of land across the EHJV. We set quantifiable objectives based on species abundance, and mathematically identified the subset of planning units which most efficiently reach those objectives. Essentially, this boils down to a minimum set optimization problem (Wilson et al. 2009): **what is the smallest number of grid cells which, combined, contain at least X**% **of the population of each priority species within the EHJV?** By solving this problem for different values of the target *X*, we produce maps which rank every grid cell according to its contribution to the optimization objective. The principal contribution of this methodology is its ability to balance multiple conservation goals. If there were only one species included, the conservation priority map would look very much like its population distribution map. The benefit offered by the optimization routine we apply lies in its ability to identify areas which contribute to multiple conservation objectives simultaneously, while also ensuring that every objective meets its assigned coverage target so that no species is left out. We opt to use uniform fractional targets across all features (i.e., to seek the same *X*% coverage for each species, expressed as a fraction of that species’ total region-wide abundance) for the sake of simplicity and impartiality. Non-uniform approaches which weight species according to some metric of ecological importance or threat status could also be considered (Joseph et al. 2009); however, in the absence of any study quantitatively deriving such weights or any priority hierarchy laid out in the EHJV Implementation Plan, we treated all target species equally.

Once we have obtained the species population data, defined our planning units, and chosen the coverage target *X*, the optimization can be set up as a mixed integer linear programming (MILP) problem and solved exactly (i.e., the resulting combination of selected planning units will be guaranteed to be the minimum number that meets the designated targets) (Beyer et al. 2016). As mathematically elegant as an exact MILP solution may be, however, it presents a limited answer to our question. With as many planning units as our study area contains, there are almost certainly a large number of near-optimal solutions (i.e., different combinations of the same number grid cells which offer almost the same population coverage as the true optimum, falling short of it by only a small fraction of a percent). For a real-world problem where every data point comes with some margin of error, all of these solutions must be treated as essentially equally plausible. We therefore compute composite solutions by averaging over an ensemble of 100 near-optimal results for each chosen coverage target. These calculations are carried out using the `prioritizr` package in R (Hanson et al. 2024), which offers the functionality of generating “portfolios” of MILP solutions in the vicinity of the true optimum. The portfolio size of 100 represents a conservatively chosen threshold: computed means are generally observed to change very little for ensembles larger than this.

The choice of this *X*% coverage target, while sometimes anchored in policy (e.g., Eckert et al. (2023); Karimi et al. (2025)), can be difficult to relate directly to overarching objectives such as preventing extinction or maintaining populations. Therefore, instead of committing to any specific target, we sweep over values from 0-100% for all species. By averaging these results, we can assign a priority score between 0 and 1 to each cell of the grid. We have restricted our analysis to problems with uniform coverage targets (i.e., the same target for each species, expressed as a fraction of that species’ total region-wide abundance). In this formulation, the priority scores represent the proportion of near-optimal solutions across the swept target range in which a given cell is selected, which can be interpreted as a measure of how high the coverage target has to be for the cell to be included in most solutions. A score of 0.7 means that that cell is contained in a majority of solutions with at least 30% coverage of all species. For a score of 0.9, it is included in a majority of solutions with at least 10% coverage. This offers a comparative evaluation of the conservation value of all grid cells in a single visual representation, as shown in Fig. 1.

**Fig. 1:**
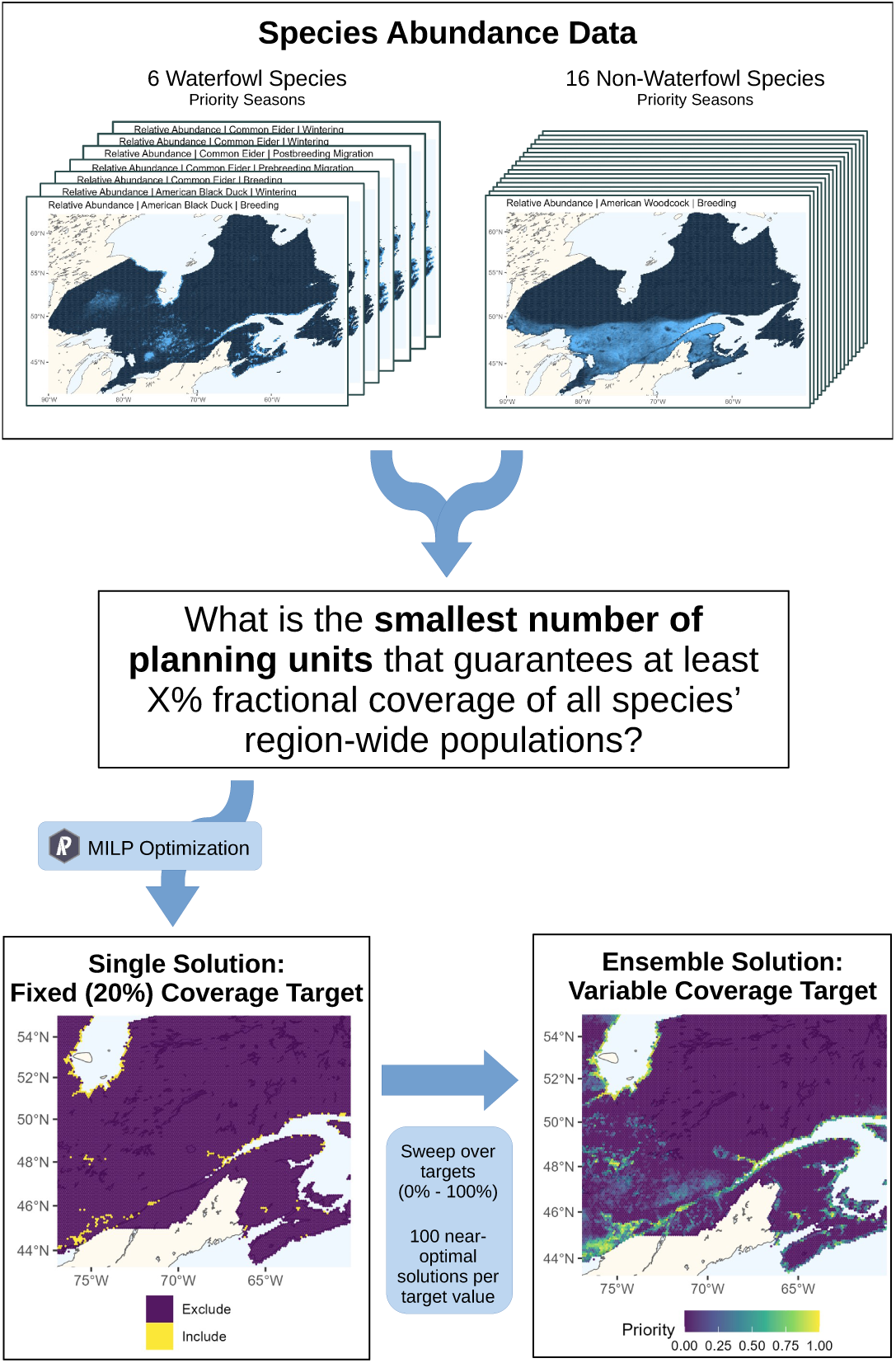
Conceptual schematic for incorporating relative abundance data of 22 priority bird species to identify priority areas for conservation in Canada’s Eastern Habitat Joint Venture. The discrete MILP solutions are aggregated to obtain quasi-continuous prioritization scores on a 0-1 interval. The bottom right map can, to good approximation, be recovered from the bottom left by thresholding the priority score distribution at 0.8 (corresponding to its 20% coverage target).

### 2.4 Seasonal Data Considerations

The relative abundance data published by eBird is temporally resolved (intra-annually), so our analysis can be carried out with attention to particular times of year, as defined by each species’ breeding, migration, and wintering seasons (approximate dates for which are published with the eBird data products). Unsurprisingly, the geographic distributions of the species in question vary considerably across these seasons. One logical method for synthesizing these results into a single annual prioritization map would be to perform a single optimization across all seasons, weighted by their durations (Schuster et al. 2019). The shortcoming of this approach is that it treats each day of occupancy as equally important to the species’ life cycle. One common alternative is to focus only on protecting breeding habitats, which are the most crucial for waterfowl population growth (Hoekman et al. 2002; Sedinger et al. 2019), but some have criticized this as an overly narrow framing (Marra et al. 2015; Runge et al. 2015). We considered seasons on a species-by-species basis (Table 1) according to the priorities laid out in the EHJV’s species white papers (Eastern Habitat Joint Venture Science Team 2026). Thus, our conservation objectives included coverage goals for 34 species/season combinations. Species represented in more seasons may have more influence on solutions, depending on the degree of overlap across their seasonal abundance distributions (e.g., if the coverage required to meet a species’ breeding season target also provides sufficient coverage to meet its migratory season targets, then the inclusion of those additional seasons will not affect the optimization results). Prioritization results based on alternate formulations, including season-by-season results and year-round results, are included in Appendix B.

**Table 1:**
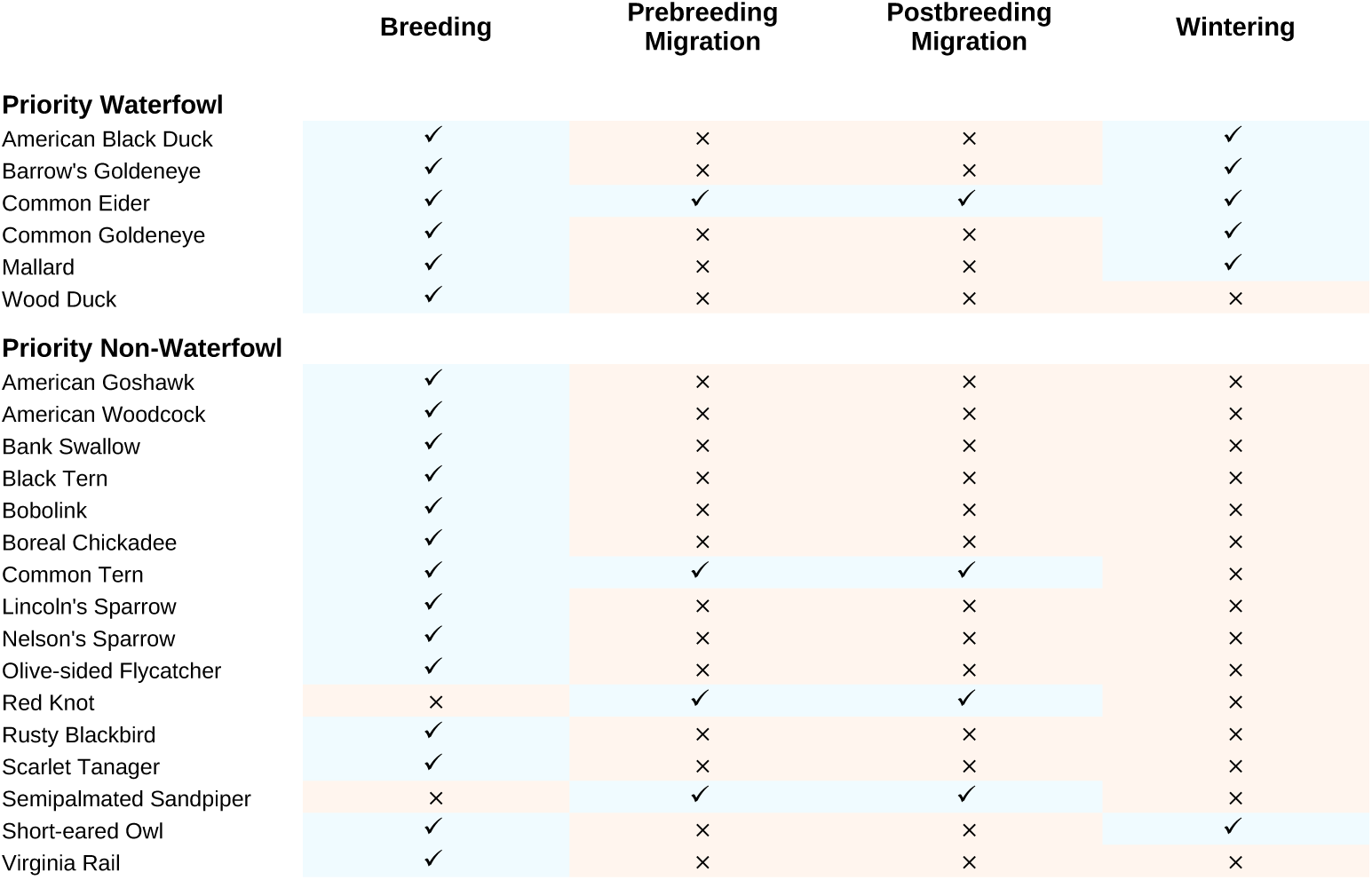
Seasonal abundance data inclusions in the primary prioritization results for each priority species in Canada’s Eastern Habitat Joint Venture.

### 2.5 Adding Geographical Constraints

The prioritization approach we have outlined estimates value in purely ecological terms, without consideration for the mode or regional scope of the conservation measures whose deployment it might help to inform. This is a design choice intended to maximize the generality and interpretability of the results. We recognize, however, that a comparative evaluation across the full span of the EHJV without regard for land ownership status or political jurisdiction may not meet an end user’s practical needs. One important dimension to consider is the incorporation of knowledge of existing protected areas, as well as areas which fall outside the practical jurisdiction of most conservation action. Based on the Canadian Protected and Conserved Areas Database (CPCAD), we estimate that approximately 12% of the area of the EHJV already has protected status (Government of Canada; Environment and Climate Change Canada; Canadian Wildlife Service 2025), though this figure may be higher, as CPCAD does not include all protected areas. If one priority species has a large amount of habitat already within these areas while another does not, it stands to reason that a prioritization scheme for future conservation planning should give greater attention to the latter. We integrate this principle into our optimization scheme using the maps published by CPCAD: any grid cell which is at least 50% covered by protected land is included by default in the algorithm’s solution, and the species abundance values associated with that cell are counted toward the designated coverage targets. As a result, the remaining (unprotected) grid cells that the algorithm selects are those which most efficiently close the gap between the existing coverage and the coverage targets. This 50% coverage threshold means that some grid cells labeled on our map as “protected” may contain some minority area of non-protected land. These cells are included in solutions by default and thus will not be assigned prioritization scores by our methodology, but this does not mean that those areas are of low conservation value (if anything, a grid cell already dominated by land deemed worthy of protection under existing designations is more likely to have high ecological value).

Additionally, some users may wish to exclude grid cells which may not be eligible for conservation action. While we do not omit any grid cells on this basis when computing prioritizations, we provide maps with shaded overlays to indicate potentially unavailable lands in Appendix F. Two possible exclusion classes are plotted: one illustrating lands held in reserve by the Federal government for military or other purposes, based on data from the Canadian Forest Service’s Map of Forest Management in Canada (Government of Canada; Natural Resources Canada 2020), and one indicating Indigenous lands as demarcated by Natural Resources Canada’s Aboriginal Lands of Canada Legislative Boundaries map (Government of Canada; Natural Resources Canada 2024). Our choice to not exclude Indigenous lands from our primary analysis does not imply availability of these lands for top-down federal or provincial conservation measures by government agencies or NGOs. Rather, it is a choice not to exclude Indigenous lands from potential Indigenous-led or co-led initiatives to help achieve mutually-beneficial goals.

Lastly, while results computed across the full area of the EHJV are likely to be useful for a conservation planner who wishes to make comparative evaluations of delivery sites across the entire region, they may be less useful for planners operating within smaller jurisdictions. Considerable conservation planning in the joint venture occurs at the provincial level (and often more locally), and results computed EHJV-wide may not be useful for planning at this scale. To address this shortcoming, we present alternative maps based on computations carried out for each province individually (i.e., species coverage targets are calculated as fractions of each species’ abundance within that province). We have included these maps in Appendix C. The same method is applied in Appendix D to produce separate maps for publicly- and privately-owned lands, which often require different approaches for conservation delivery. This frame-work assures that sites of high regional conservation value are highlighted in every jurisdiction. It should be noted, however, that the solutions define different effective scales for each region—a priority score of 0.8 in Quebec is not equivalent to a score of 0.8 in Nova Scotia, in terms of the absolute species abundance coverage it denotes.

## 3 Results

We identified relative priority scores for waterfowl and non-waterfowl species across the entire study area. The waterfowl map (Fig. 2) identifies high-priority areas along much of the marine coastline of the EHJV and the St. Lawrence River, as well as inland areas primarily in southern and western Ontario. The non-waterfowl map (Fig. 3) identifies large inland regions of high value, with particular emphasis on the Gaspé Peninsula and St. Lawrence estuary, northern Ontario along the Hudson Bay coast, central Quebec, and Newfoundland. Although the latter map shows a higher proportion of high-scoring grid cells than the former, this should not be understood to reflect an overall difference in priority between the two species groups. Rather, the more extensive spatial coverage on the non-waterfowl map results from 1) a larger number of species with coverage targets to meet, and 2) comparatively high spatial dispersal in non-waterfowl versus waterfowl. The two maps result from separate computations and are plotted on two (internally consistent but incommensurate) color scales.

**Fig. 2:**
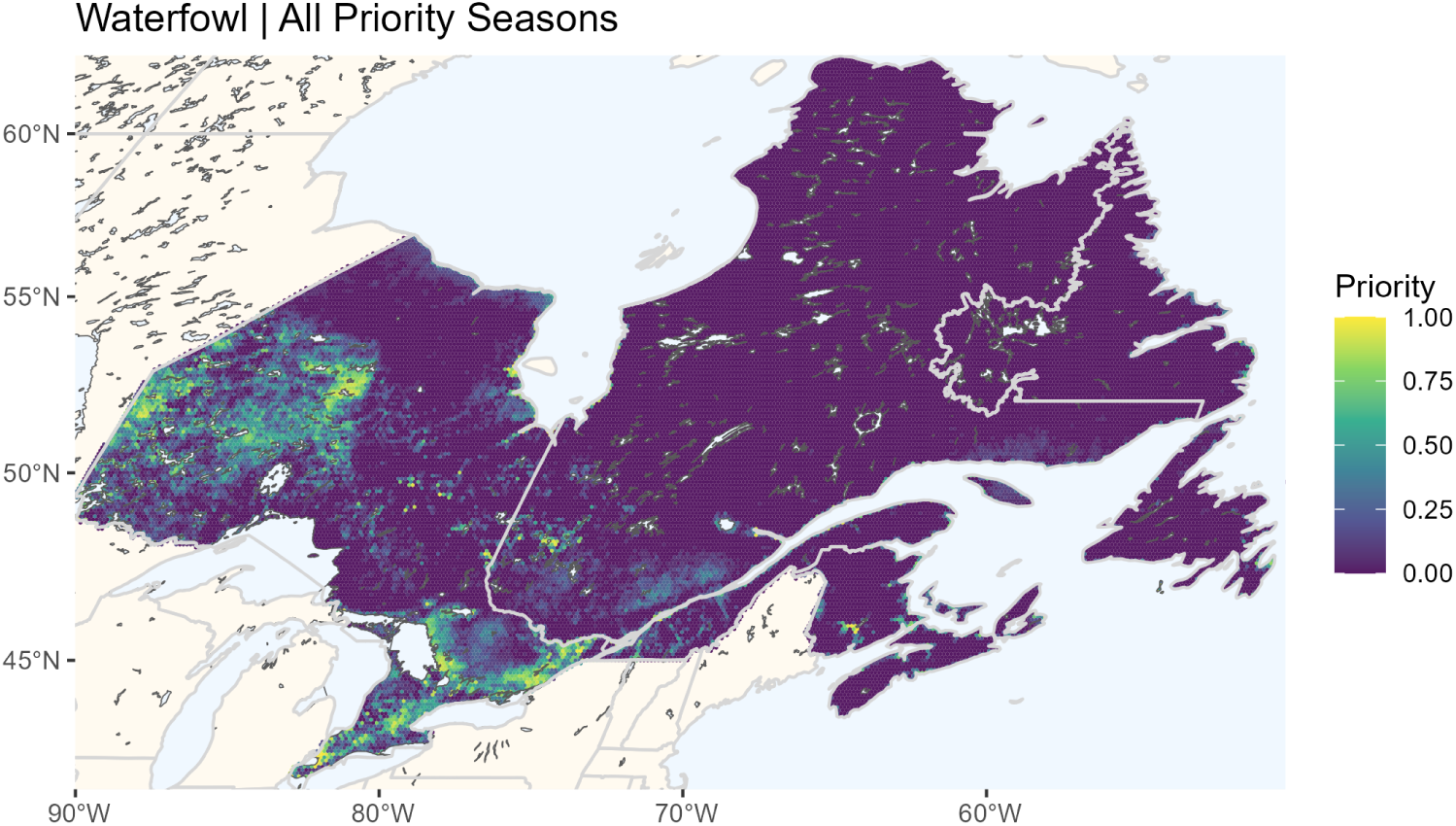
Prioritization map for the 6 priority waterfowl species within Canada’s Eastern Habitat Join Venture, based on their distributions during the seasons specified in Table 1.

**Fig. 3:**
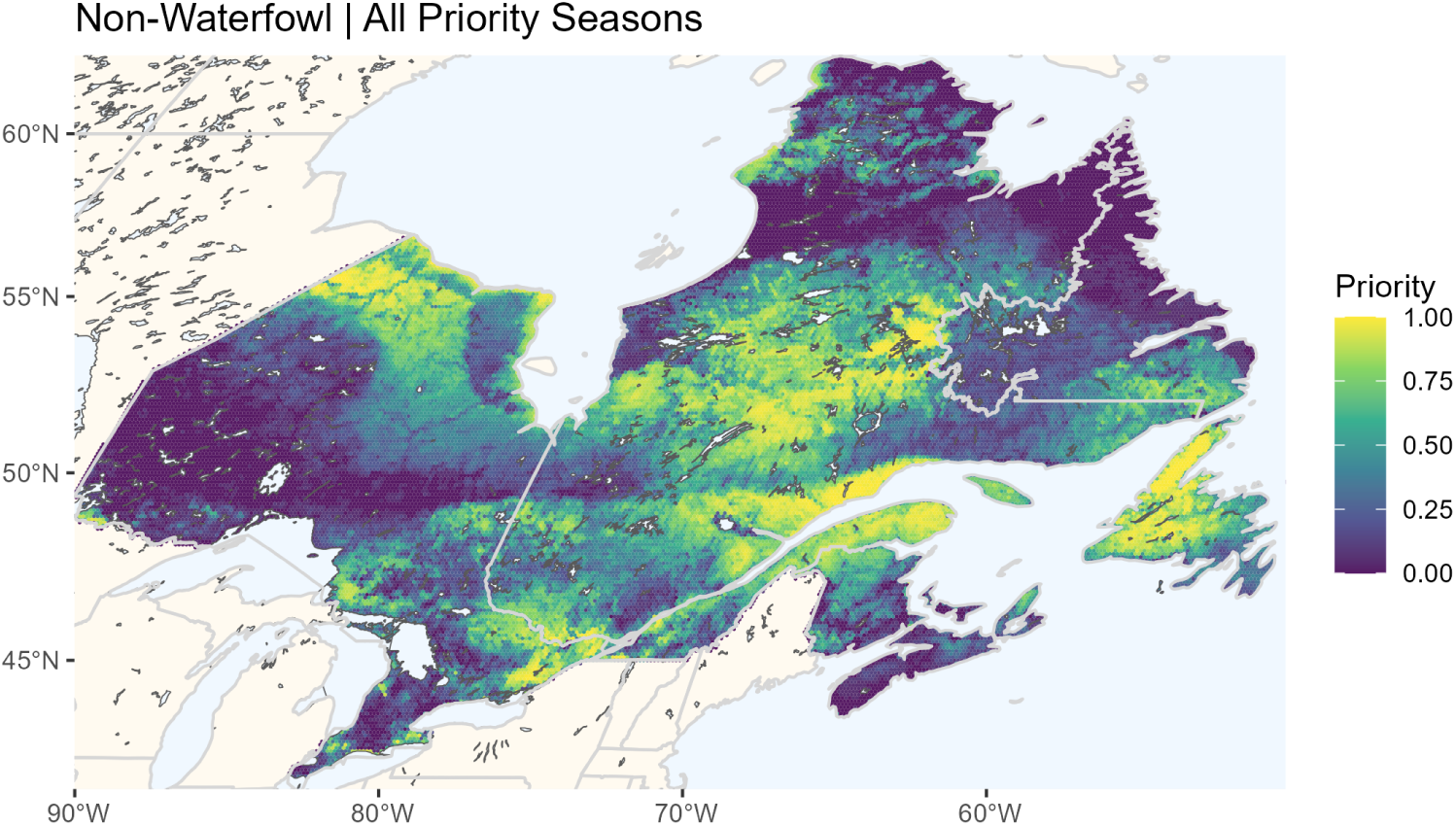
Prioritization map for the 16 priority non-waterfowl species within Canada’s Eastern Habitat Join Venture, based on their distributions during the seasons specified in Table 1.

### 3.1 Geographically Constrainted Prioritizations

Using data from the 6 priority waterfowl species during their included seasons, we compared results obtained with and without designating existing protected areas as “locked in” to solutions. Prioritization maps derived with and without these constraints are presented in Fig. 4 for comparison.

**Fig. 4:**
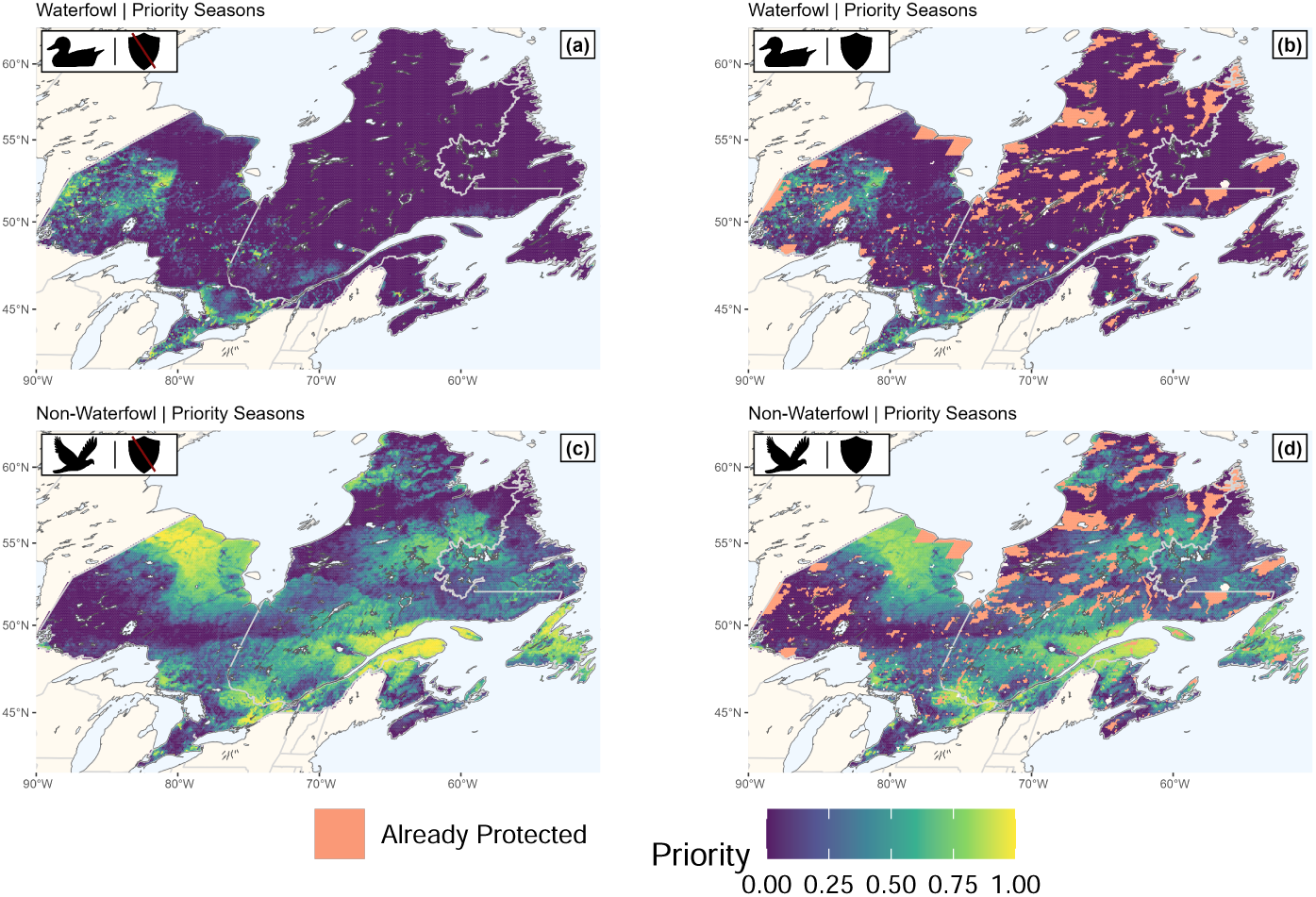
Prioritization maps for priority waterfowl (top) and non-waterfowl (bottom) species within Canada’s Eastern Habitat Join Venture. Left: prioritization schemes for the full EHJV area. Right: prioritization schemes with existing protected regions (pale yellow) locked in to solutions by default.

Our focal waterfowl species varied in the proportion of their populations already covered by existing protected areas from *<* 5% to about 30% (illustrated on the y-intercept of Fig. 5). The highest priority values (the top *∼* 4, 000 km^2^ of the study region) are assigned preferentially to cells with high Wood Duck (*Aix sponsa*) abundance, as that is the species with the lowest fractional coverage. As this gap is closed, other species are targeted in turn until their fractional coverages have been equalized, and further cells are prioritized with equal attention to all species. The exception to this is the Barrow’s Goldeneye (*Bucephala islandica*), whose small population in the EHJV is already disproportionately covered by protected areas. Incidental increases in its coverage from areas prioritized for the other species’ benefit, particularly that of the often co-located Common Goldeneye (*Bucephala clangula*), are sufficient to ensure that the resulting prioritization scheme is more than sufficient for it to meet and even exceed the same fractional target as the other species. Only the highest priority scores are affected by differences in species’ existing coverage. The disparities resulting from these different starting points are only reflected in the top *∼* 15, 000 km^2^ of scores, which represents less than 0.5% of the total EHJV area (about 2.8 million km^2^). As a result, accounting for existing protected areas has a proportionally small impact on prioritization distributions (beyond the fact that those regions are themselves omitted from solutions).

**Fig. 5:**
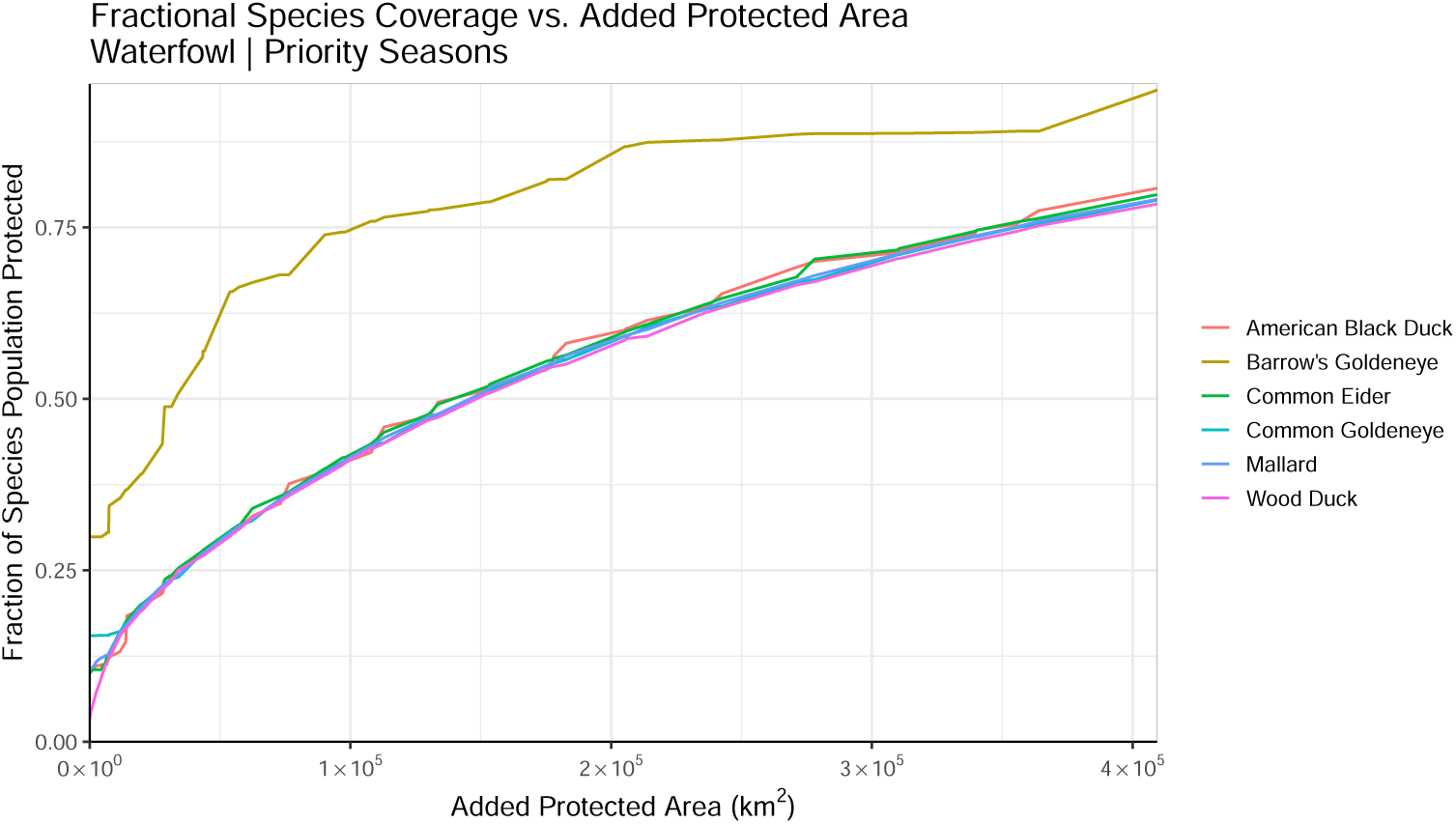
Fractional coverage of each priority waterfowl species in Canada’s Eastern Habitat Join Venture, as new protected areas are added (in order of assigned priority score).

## 4 Discussion

Flexible and transparent decision-support tools that integrate the best available data and can be readily updated are essential for effective conservation planning. Our ensemble-based, or sweep, approach applied here provides a generalized assessment of conservation value across the entire EHJV. Rather than a binary solution which reflects a single, fixed coverage goal or area budget, it offers spatially explicit insight for conservation practitioners operating at a range of spatial scales. For the EHJV, the prioritization surfaces provide a contemporary representation of the areas most likely to contribute toward achieving population objectives while minimizing the area required to do so. Even in the absence of explicit relationships between habitat program actions and population responses, this framework helps decision makers identify where conservation investments have the greatest potential to benefit priority populations.

Our objective was not only to produce an updated prioritization product, but to demonstrate a transparent and repeatable workflow capable of generating and refining such products through structured decision-making and stakeholder engagement. Conservation priorities, available data, and management objectives will inevitably evolve, and the framework is designed to evolve alongside them. We identify numerous opportunities for expert knowledge to be formally incorporated into the process, including decisions regarding species selection, species weighting, seasonal periods, and the treatment of species individually or collectively within prioritization analyses. These choices can substantially influence prioritization outcomes and reflect important trade-offs among conservation objectives, species distributions, and opportunities for conservation delivery.

### 4.1 Abundance Data Limitations

We used eBird abundance layers as our principal data sources primarily based on their species coverage, spatial extent, and seasonal resolution. We emphasize that this decision should not be interpreted as an endorsement of their accuracy relative to other models. Indeed, their use of user-submitted checklists introduces potential sources of bias and measurement error as compared to the other candidate products, which are primarily derived from more standardized aerial survey data (Frei et al. 2024; Adde et al. 2020a). The relative merits of these data collection approaches for the purpose of abundance modeling are the subject of ongoing debate (Robinson et al. 2025; Shen et al. 2023; Feng and Che-Castaldo 2021; Winiarski et al. 2024). Aerial surveys are designed to quantify indicated breeding pairs and typically exclude large non-breeding flocks (Frei et al. 2024), whereas eBird records all birds present regardless of breeding status (Fink et al. 2025). As a result, locations with high eBird-derived scores may reflect staging or non-breeding aggregations rather than breeding habitat value. While no data products other than the eBird Status and Trends abundance data currently offer sufficient species coverage to constitute a viable alternative for the purposes of this study, future survey-based models may allow for more rigorous comparison of how alternate measurement methodologies affect the resulting prioritization maps. As a point of comparison, maps produced using survey-based approaches are included in Appendix E.

### 4.2 Accommodating Changing Species Distributions

Our analysis was based on static data products intended to reflect present-day conditions—the principal computational inputs come from the 2023 release of the eBird Status product (the most recent available at time of writing). However, the use of current species abundance data as a proxy for conservation value is an imperfect heuristic. While it is generally accepted that areas where a priority species congregates should be considered worthwhile targets for conservation, there may be other, less populated areas which serve as valuable *potential* habitat, subject to changes in aspects such as distributional change or habitat restoration (e.g., (Wilson et al. 2011; Possingham et al. 2015)). We make no attempt to account for this latter category in our analysis because identifying such locations entails significant conjecture and cannot easily be integrated into our quantitative framework.

Likewise, we do not endeavor to project future changes to species abundance due to changes in climate and land use. Using species distribution models (SDMs) to make predictions in this manner is a well-explored area (Araujo and New 2007; Araújo et al. 2005; Elith and Leathwick 2009; Franklin 2010, 2013; Schuster et al. 2023; Adde et al. 2020b), but it does introduce a number of additional sources of error (Peterson et al. 2018). In addition to the uncertainties which inevitably accompany the projections of input variables, there is no guarantee that the SDM itself will be robust to extrapolation: these models are derived from correlative relationships between environmental covariates and species occurrence, but those associations cannot necessarily be assumed to persist under new conditions which may differ from those in their empirical training data. We have elected to avoid the added challenges associated with using future-state projections of species distributions. The EHJV implementation goals guiding our analysis set a planning horizon of 2030, which is in line with the existing data products (circa 2023). We do feel that this would be a valuable line of future inquiry, particularly as a resource for long-term or climate-sensitive conservation planning.

Our choice to take a static approach means that the prioritization schemes presented represent snapshots based on current population distributions—valuable for conservation planning under present conditions, but perhaps less well suited for development of long-term plans which involve substantial reconfiguration of populations. Crucially, this does not generally apply to small-scale restoration efforts: for restoration of habitat on a scale significantly smaller than our planning units, unit-wide species abundance still serves as a useful proxy for a project’s value. We expect that this assumption weakens with increasing intervention size; larger interventions which lead to significant, localized population changes at a scale approaching (or above) the grid scale may fall outside the scope of our planning framework. The size of our chosen planning units (*∼* 87 km^2^) is intended to significantly exceed the typical scale of restoration intervention projects in order to mitigate this effect.

### 4.3 Incorporating Additional Conservation Values

Our current prioritization framework prioritizes areas based solely on the abundance distributions of the focal species. However, there exist many candidate covariates related to sociopolitical interests, ecosystem services, land-use pressures, or stakeholder investments which could feasibly be leveraged to broaden the scope of our results. The incorporation of many of these factors would likely involve decisions tailored to specific conservation goals or delivery mechanisms, which is outside the scope of our current study. Nonetheless, we recognize that many on-the-ground implementation measures could benefit from the inclusion of additional inputs, so we offer here a brief discussion of how such values might be integrated.

In some cases, additional factors may be suitable for direct inclusion in the optimization framework. This could be accomplished by maximizing their value alongside that of species abundance (Joseph et al. 2009), using them to constrain solutions or weight planning units (Jung et al. 2021), or modifying the formulation of the objective function (Daigle et al. 2020; Hanson et al. 2022)). For example, flood-risk information could be used to identify areas where conservation provides additional co-benefits such as flood attenuation. This would require determining whether flood-prone areas should be upweighted (to support co-benefit delivery) or downweighted (to avoid high-risk investments), depending on partner objectives.

Alternatively, non-biological values can remain external to the optimization, but still be provided as supplemental geospatial layers to inform partner decision-making. This approach preserves the interpretability and transparency of the core biological prioritization while still enabling partners to apply other decision criteria relevant to their region, mandate, or community. We recommend prioritizing based on the abundance of EHJV priority species, in line with the Joint Venture and NAWMP objectives, and using additional values as supplemental information that may support conservation decisions and funding requests by partners.

## 5 Conclusion

The prioritization schemes presented in this study represent the best available recommendations for conservation planners seeking to identify areas that contribute maximally to population coverage across all included species. These results are mathematically optimal with respect to this objective and the available data, but we emphasize that they should not be interpreted as unilateral prescriptions for on-the-ground conservation action. The number of birds enclosed by a given region does not solely account for the value that land offers to the species population, and even if it did, the best available species abundance models would still be imperfect representations for the true, dynamic distributions of these populations. We have endeavored to provide prioritizations tailored to the needs of practitioners planning conservation action in a variety of modes and geographical scopes, but in all cases these products should be considered alongside existing knowledge of local ecosystems and the specific conservation goals of the user. The quantitative, top-down approach of our analysis is designed to highlight opportunities which may not be obvious to the human eye—areas which might not hold notable value to any single priority species, but which emerge as strong conservation candidates by offering moderate benefit to multiple objectives simultaneously. In this respect, the results of this study constitute an instrumental decision-making resource to support the collective conservation of all priority species within the EHJV.

## 6 Acknowledgments

This work was completed with financial support from the Black Duck Joint Venture (United States Fish and Wildlife Service; USFWS), a Mitacs Accelerate fellowship to DD, and Environment and Climate Change Canada (Government of Canada). We are grateful for feedback from the EHJV Management Board and Science Committee, which reviewed earlier draft results. The content is solely the responsibility of the authors and does not represent the official views of the USFWS, the Government of Canada, or of the EHJV.

