## Supplemental Information for "Optimal Conservation Prioritization for Waterfowl and Other Birds in Eastern Canada"

### Appendix A Code and Data Availability

The species abundance data used in this analysis is available through the eBird Status and Trends (Fink et al. 2025) web portal at https://science.ebird.org/status-and-trends/download-data. Additional abundance data used to generate alternative results in Appendix E were generated by Frei et. al. (2024) (Frei et al. 2024) and provided to us by the authors of that study. Other data products used to support this analysis are made available by the Government of Canada. The Canadian Protected and Conserved Areas Database (CPCAD) is available at https://open.canada.ca/data/en/dataset/6c343726-1e92-451a-876a-76e17d398a1c. The Aboriginal Lands of Canada Legislative Boundaries data product is available at https://open.canada.ca/data/en/dataset/522b07b9-78e2-4819-b736-ad9208eb1067. The Canadian Forest Service's Map of Forest Management in Canada is available at https://open.canada.ca/data/en/dataset/d8fa9a38-c4df-442a-8319-9bbcbdc29060.

R code to reproduce the results presented in this paper is available on GitHub at https://github.com/dylewsky/Optimal_Conservation_Prioritization. This code provides minimal examples to preprocess raster abundance data (from eBird or otherwise) and generate prioritization maps similar to those published in this study.

In addition to the static prioritization maps presented in this document, we have designed our data processing pipeline to be compatible with the Where to Work tool (Hanson et al. 2026) managed by the Nature Conservancy of Canada. This interface, available as a web app at https://ncc-cnc.github.io/wheretowork/index.html, allows user to set up and evaluate optimization problems using the same underlying methodology used in this report (implemented through the `prioritizr` package in R). To support this compatibility, our preprocessing pipeline is a lightly modified version of code published by the Where to Work team at https://github.com/NCC-CNC/wtw-data-prep/, reproduced in our repository with the permission of the original authors.

The compatibility of this tool is intended to offer users the ability to exercise more fine-tuned control over the prioritization scheme. It can be used to generate solutions with manually specified coverage targets for every species and lifecycle season, so one can explore specific scenarios not represented in our main corpus of results. We anticipate that most users will not require this level of customization, however, and suggest that this tool be used as a secondary resource to the already prepared prioritization maps. The Where to Work tool does not support the ensemble approach we have presented in this paper. However, by manually adjusting coverage targets, one can roughly reproduce the information conveyed in the gradated representations we have presented.

### Appendix B Season-by-Season Prioritization Results

Prioritizations developed using data from each seasonal category separately are presented in Figs. B1 and B2.

Alternative prioritizations in which all seasons are included for all species are presented in Fig. B4. Here, seasons are weighted by their durations so that results reflect

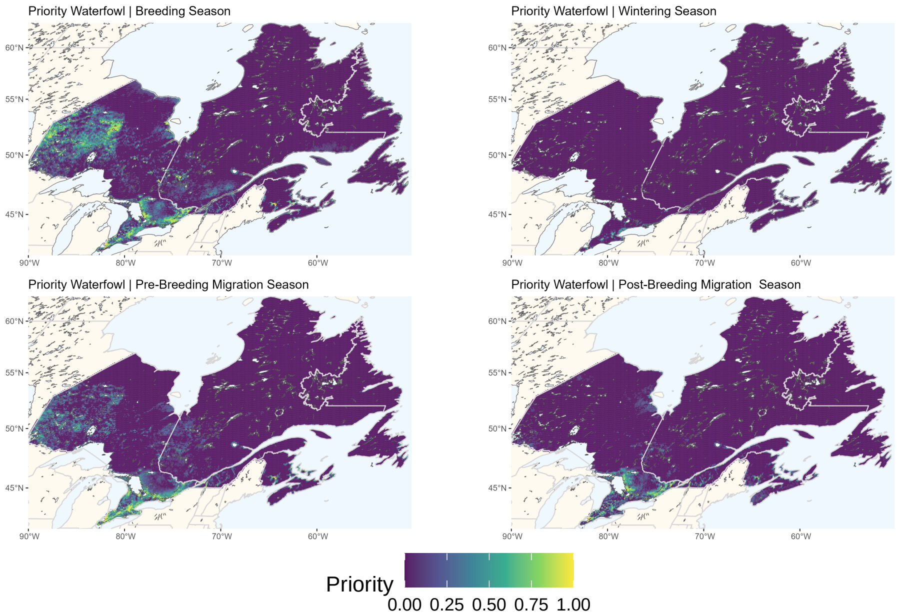


**Fig. B1**: Prioritization maps for the 6 priority waterfowl species, separated by season.

an occupancy-only valuation approach, without regard for life cycle stage. Each priority species has its own dates demarcating the seasons, but on average the migratory periods are the longest, as shown in Fig. B3.

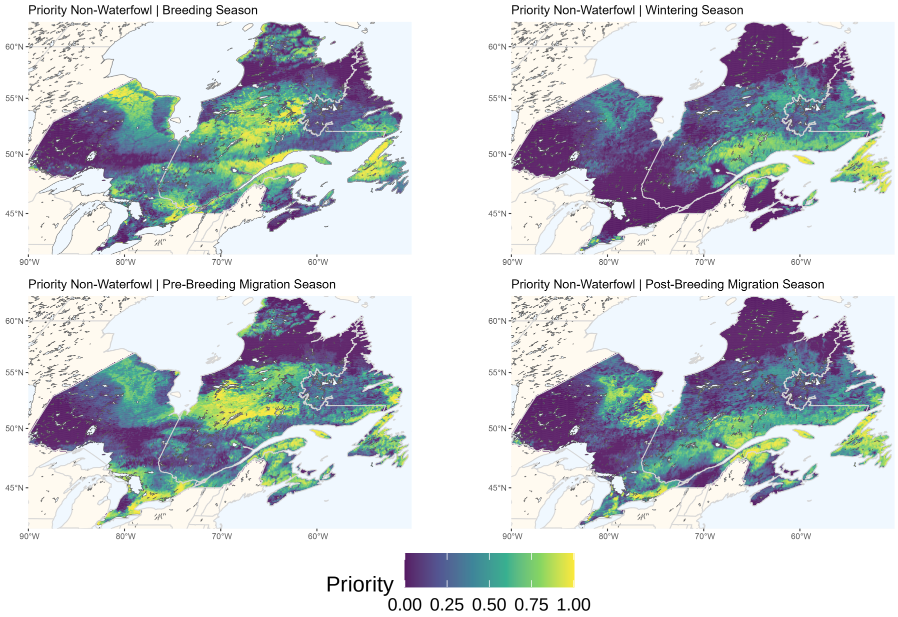


**Fig. B2**: Prioritization maps for the 16 priority non-waterfowl species, separated by season. (Note: the larger high-priority regions on these maps, as compared to those for the waterfowl species, result from the properties of the input data. There are more priority non-waterfowl species, and their populations tend to be more geographically dispersed. Hitting a given coverage target (e.g. 20% of each species' population) therefore necessitates a larger solution area for this data set than the same target would for the waterfowl species).

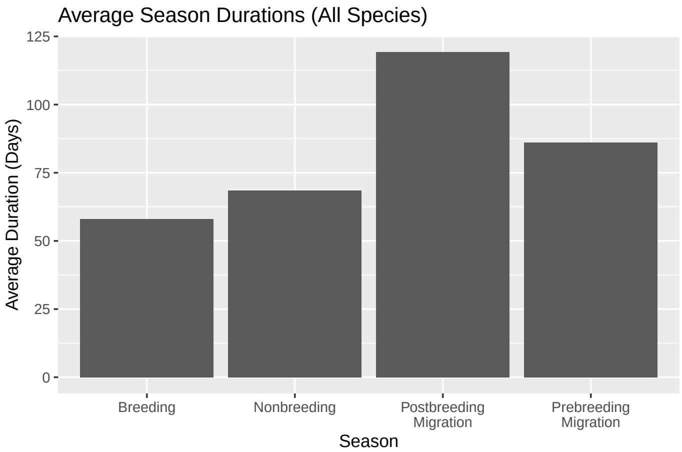


**Fig. B3**: Average durations of the four seasonal life cycle stages associated with each species. These are computed using the seasonal dates published with the eBird data, and averaged across all EHJV priority species (except the Boreal Chickadee, which is generally non-migratory).

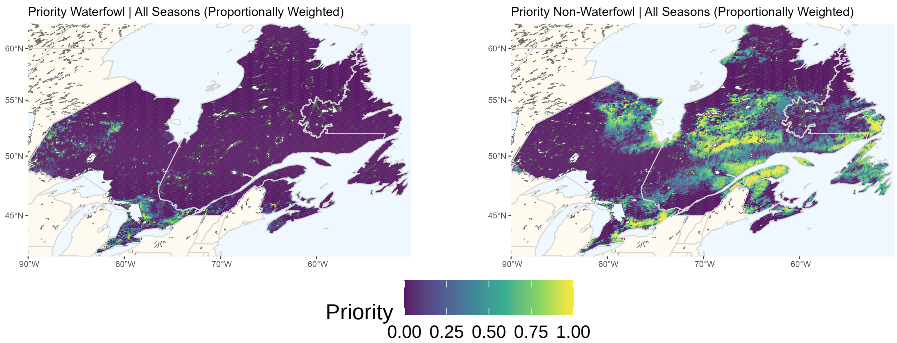


**Fig. B4**: Prioritization maps across all four seasons (breeding, wintering, pre-breeding migration, post-breeding migration), weighted for each species according to the relative duration of each season (for that species). Left: results for the 6 priority waterfowl species. Right: results for the 16 priority non-waterfowl species.

### Appendix C Separate Provincial Solutions

Separate prioritizations for each province in the EHJV are presented in Figs. C5-C9. The optimizations carried out to produce these maps treat each region individually: coverage for each species is computed as a fraction of total abundance *within that province*. Any species/season combinations with no representation (zero abundance values) across the entire province is removed from the feature pool.

Note that because the assigned priority scores are derived from fractional coverage targets set for each species, the color scales of these maps are not commensurate across provinces: a cell with a particular score on one map should not be interpreted to have equal conservation value to one with the same score on a different map. The results in this section are intended for use in intra-provincial planning only.

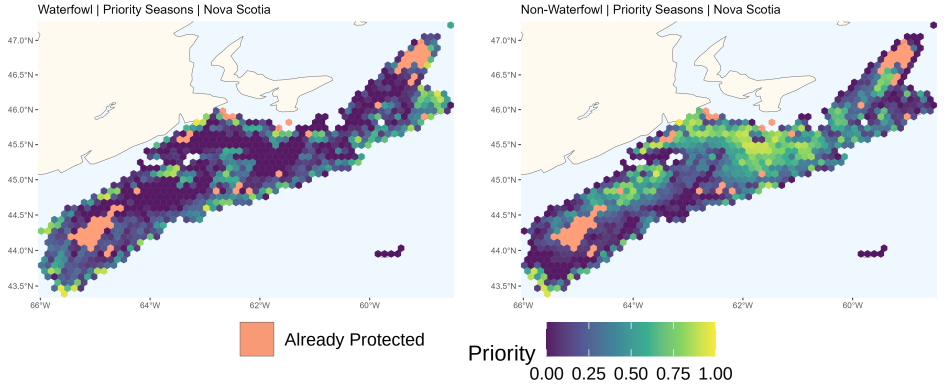


**Fig. C5**: Prioritization maps for priority waterfowl (left) and non-waterfowl (right) species during their included seasons, computed separately for the province of Nova Scotia.

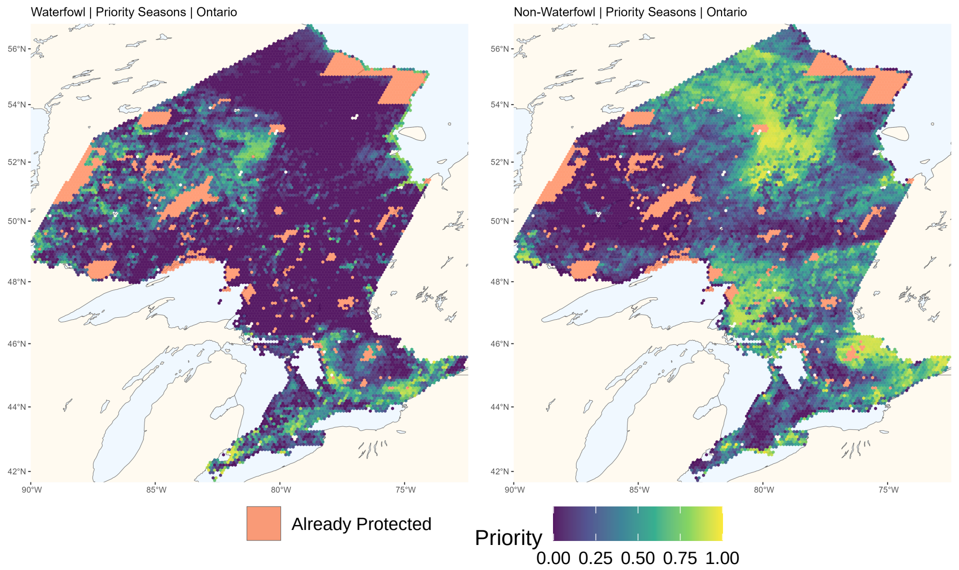


**Fig. C6**: Prioritization maps for priority waterfowl (left) and non-waterfowl (right) species during their included seasons, computed separately for the province of Ontario.

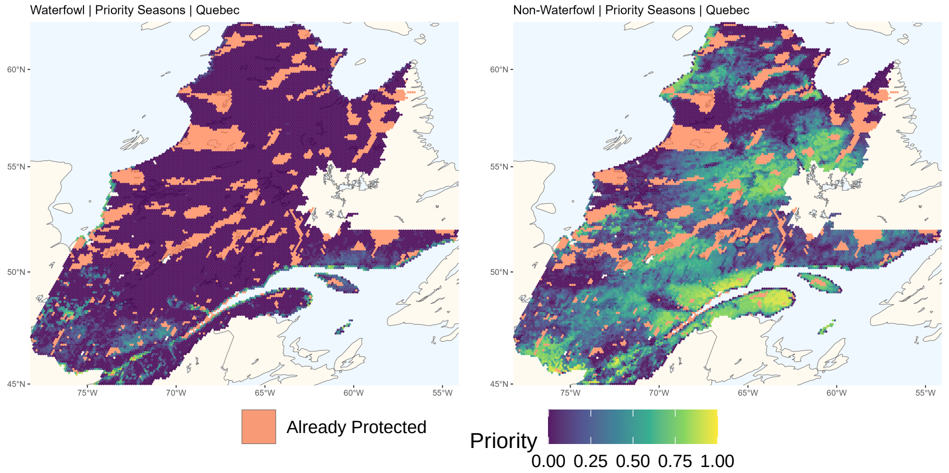


**Fig. C7**: Prioritization maps for priority waterfowl (left) and non-waterfowl (right) species during their included seasons, computed separately for the province of Quebec.

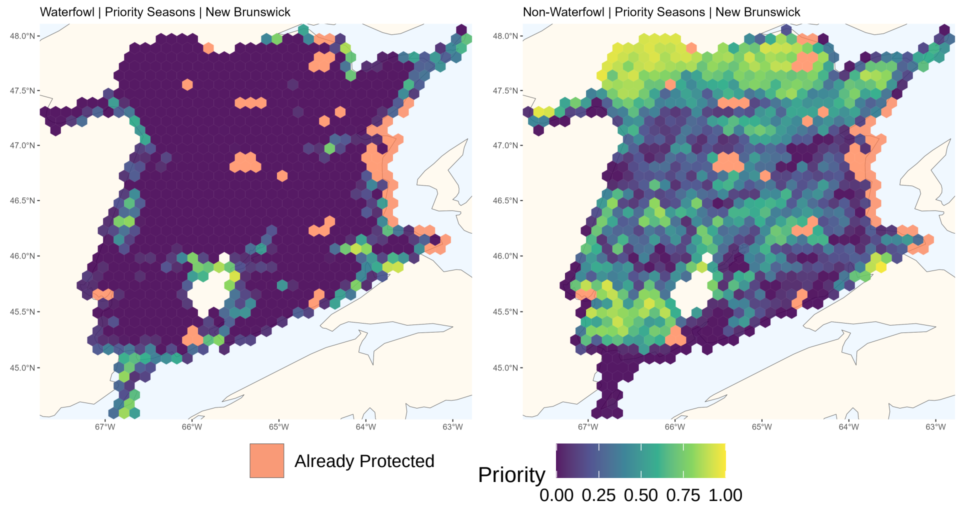


**Fig. C8**: Prioritization maps for priority waterfowl (left) and non-waterfowl (right) species during their included seasons, computed separately for the province of New Brunswick.

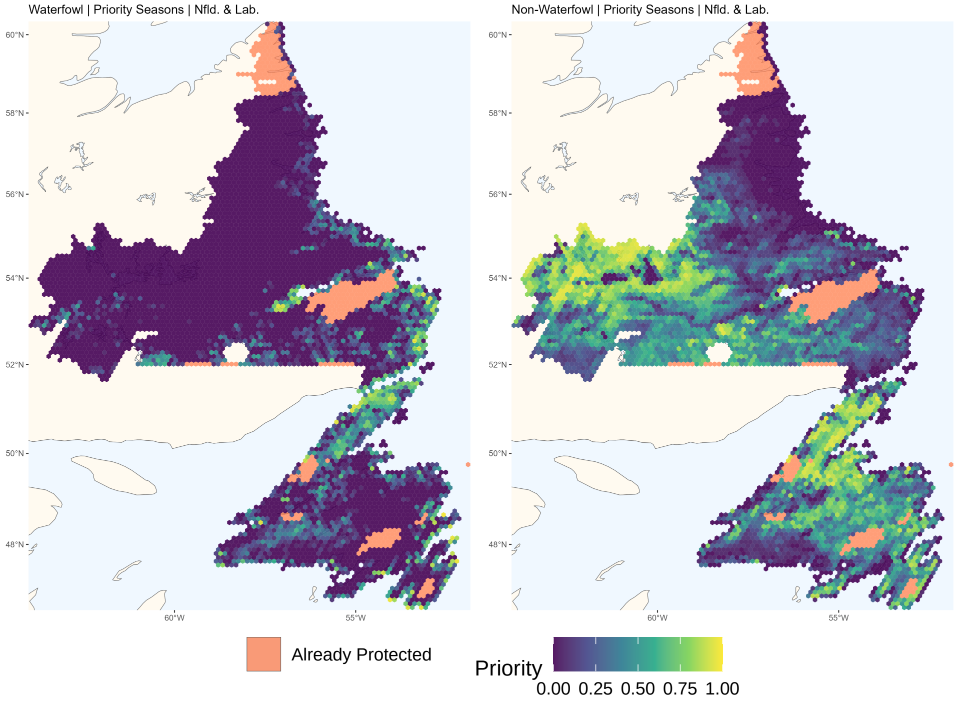


**Fig. C9**: Prioritization maps for priority waterfowl (left) and non-waterfowl (right) species during their included seasons, computed separately for the province of Newfoundland and Labrador.

### Appendix D Separate Maps For Private vs. Public Land

Similar to the results in Sec. C designed for work carried out at the provincial level, we present here an alternative formulation based on another important jurisdictional distinction: that of private lands versus public (i.e., Crown) land. Privately owned land is plotted from the Map of Forest Management in Canada published by the Canadian Forest Service (Government of Canada; Natural Resources Canada 2020). All grid cells with at least 10% private land coverage are designated as private, under the assumption that the availability of even a fraction of a grid cell for acquisition or stewardship is worthy of planning attention. All other grid cells are grouped together as "non-private," a category consisting primarily of Crown land. The cell classifications of our planning grid are shown in Fig. D10. Considering species populations for each of these two geographic subsets separately, we obtain the prioritizations shown in Fig. D11.

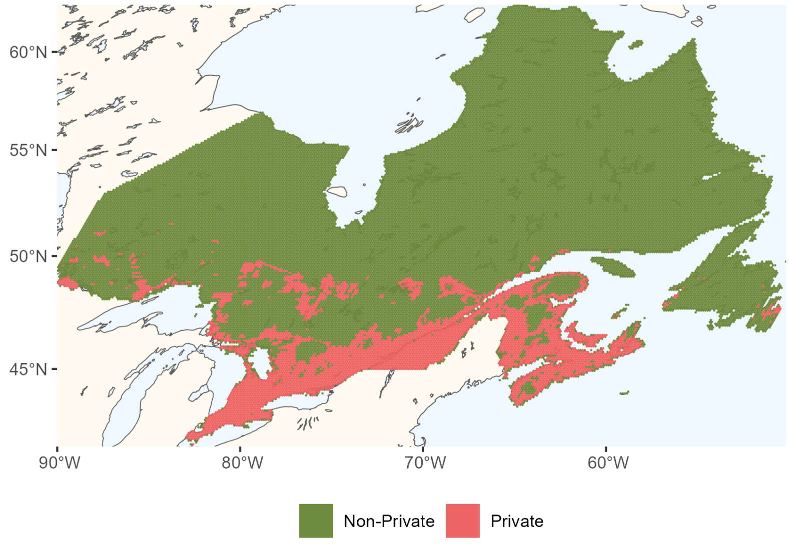


(a)

**Fig. D10**: Classification of each hexagonal grid cell labeled according to how much privately-owned land it contains.

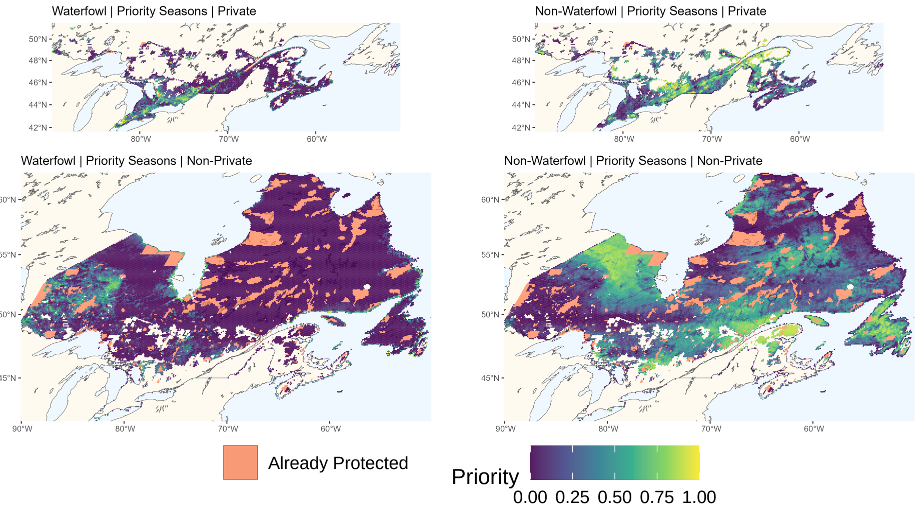


**Fig. D11**: Prioritization maps for priority waterfowl (left) and non-waterfowl (right) species during their included seasons, computed separately for regions with at least 10% private ownership (top) and primarily public lands (bottom).

### Appendix E Alternative Abundance Data Results

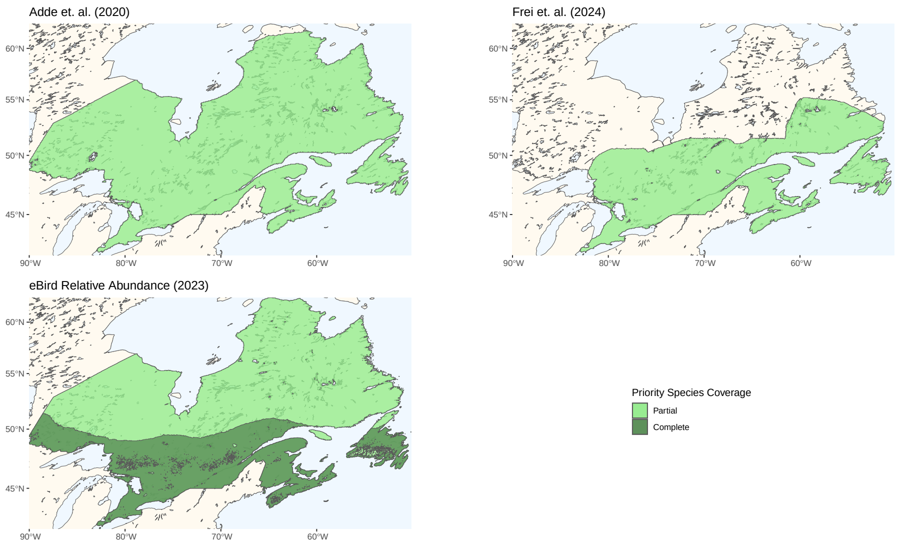


**Fig. E12**: Comparison of three candidate sources of species abundance data (Frei et al. (2024); Adde et al. (2020); Fink et al. (2025)) based on the spatial coverage they offer for some or all of the EHJV priority species.

**Table E1**: Comparison across available sources of georeferenced abundance data for EHJV priority species.

| | Frei et. al<br>2024 | Adde et. al.<br>2020 | BAM Generalized National Models v4<br>2020 | eBird Relative Abundance<br>2023 |
|---|---|---|---|---|
| **Priority Waterfowl** | | | | |
| American Black Duck | ✓ | ✓ | × | ✓ |
| Barrow's Goldeneye | × | × | × | ✓ |
| Common Eider | × | × | × | ✓ |
| Common Goldeneye | ✓ | × | × | ✓ |
| Mallard | ✓ | ✓ | × | ✓ |
| Wood Duck | ✓ | × | × | ✓ |
| **Priority Non-Waterfowl** | | | | |
| American Goshawk | × | × | × | ✓ |
| American Woodcock | × | × | × | ✓ |
| Bank Swallow | × | × | ✓ | ✓ |
| Black Tern | × | × | × | ✓ |
| Bobolink | × | × | ✓ | ✓ |
| Boreal Chickadee | × | × | ✓ | ✓ |
| Common Tern | × | × | × | ✓ |
| Lincoln's Sparrow | × | × | ✓ | ✓ |
| Nelson's Sparrow | × | × | ✓ | ✓ |
| Olive-sided Flycatcher | × | × | ✓ | ✓ |
| Red Knot | × | × | × | ✓ |
| Rusty Blackbird | × | × | ✓ | ✓ |
| Scarlet Tanager | × | × | ✓ | ✓ |
| Semipalmated Sandpiper | × | × | × | ✓ |
| Short-eared Owl | × | × | × | ✓ |
| Virginia Rail | × | × | × | ✓ |
| **Data Description** | | | | |
| Spatial Resolution | 5 km x 5 km | 5.9 km x 5.9 km | 1 km x 1 km | 3.3 km x 3.3 km |
| Model Training Data | - Eastern Waterfowl Survey<br>- Southern Ontario Waterfowl and Wetlands Plot Survey<br>- Southern Quebec Lowland Survey<br>- Maritime Agricultural Plot Survey | - Waterfowl Breeding Population and Habitat Survey<br>- eBird Citizen Science Data | - Boreal Avian Modelling Center Point Count Database<br>- Breeding Bird Survey<br>- Breeding Bird Atlas<br>- WildTrax ARU Database | - eBird Citizen Science Data |

The eBird Status and Trends abundance layers which form the basis of our prioritization analysis were selected primarily based on their superior coverage of the species (Table E1) and geographical area covered (Fig. E12) by the EHJV mandate. These data products are derived from distributed, citizen science observations, which may introduce biases and inconsistencies not present in more rigorously collected survey data. The leading survey-based candidate data source, published by Frei et. al. (2024), only covers 4 of the 6 priority waterfowl species (and none of the non-waterfowl species) during their breeding seasons on a domain spanning the southern latitudes of the EHJV. Despite its more limited scope, we feel it may be useful to present alternative prioritization results based on this product as a point of comparison (Fig. E13).

Comparison of the results from the two data sources highlights the importance of the abundance input to our analytical process. While many high priority "hotspots" are consistently identified across both sources, the overall spatial distributions of scores look substantially different. This can be attributed to a combination of differences in empirical observation data and disparate approaches to building distribution models from that data. One clear difference lies in the spatial diffuseness of reported abundances: whereas Frei et. al. report large regions of low but non-zero populations, the eBird models take a more conservative approach and report zeroes in many grid cells. Differences in the models' output distributions are shown in Fig. E14. We make no claim that either of these approaches is preferable or more accurate (indeed, both might be best suited to their respective training datasets), but only seek to convey the effects that such modeling decisions produce in downstream optimizations such as those carried out in this study.

Because the methodological differences across these models produce abundance layers with pronounced distributional disparities, we have thus far avoided any analytical approach which incorporates data from more than one source. There is a case to be made for doing so, however: a computation which uses products derived from a more rigorous survey-based methodology whenever they are available—and reverts to the crowdsourced eBird layers when they are not—would seem to offer the best of both worlds. Because our optimization formulation sets non-negotiable fractional coverage targets for each feature independently, this can be achieved without need for reweighting the data onto a single, commensurate scale. We emphasize, however, that results generated by such a "hybrid" approach must be interpreted with caution. Because of the distributional differences between the data sources, the priority areas identified are likely to be affected by the model on whose data they are primarily based. Practically speaking, features derived from the Frei et. al. (2024) model will generally produce more spatially diffuse priority areas than those based on eBird data—not because of any real, biological phenomenon, but resulting entirely from methodological differences between the source models. While this effect is also visible in the maps produced by the two data sources separately, it is much harder to identify when results are aggregated into a single map. Nonetheless, we recognize that a hybrid prioritization may be of interest to some readers, so we present a map generated by this approach in Fig. E15.

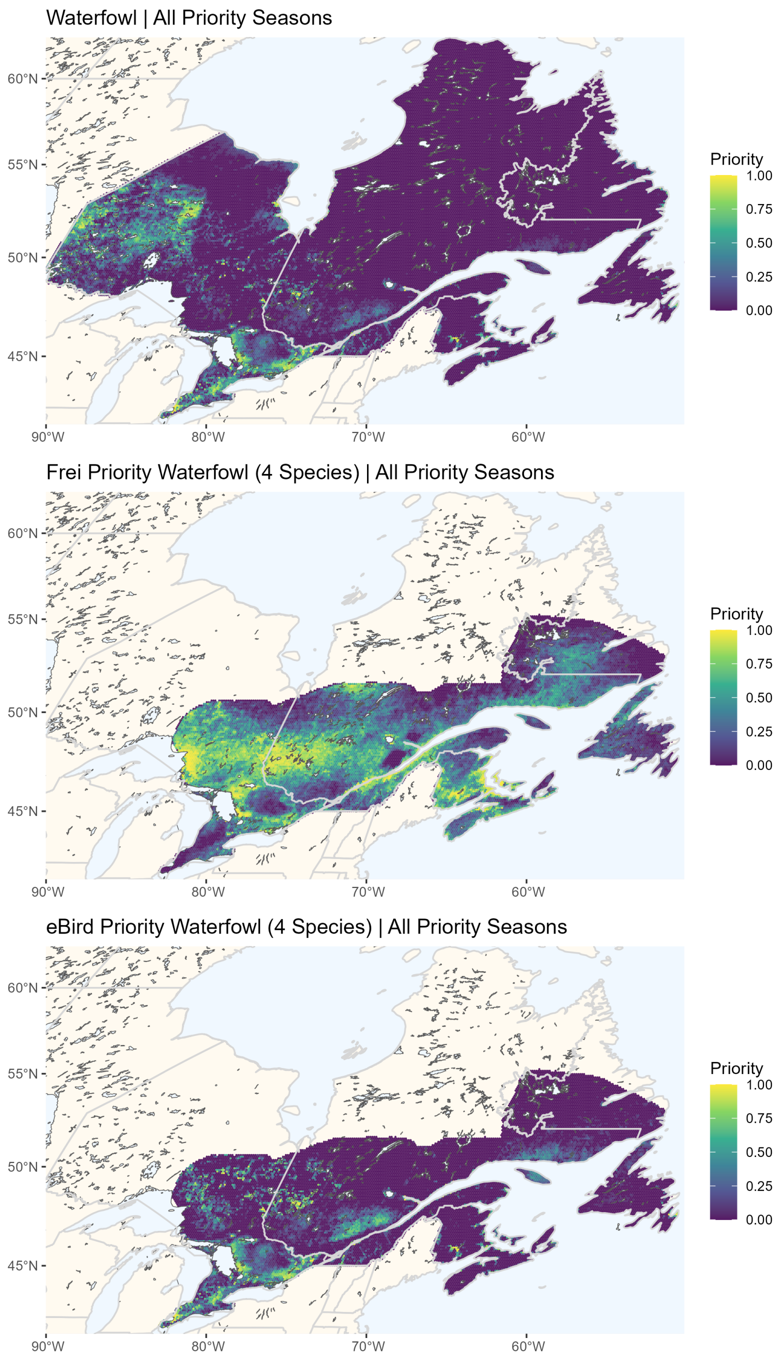


**Fig. E13**: Top: Prioritization results based on eBird abundance data across all 6 priority waterfowl species and the full EHJV region. Middle: Prioritization results based on the Frei et. al. abundance data for the 4 included waterfowl species (American Black Duck, Common Goldeneye, Mallard, and Wood Duck). Bottom: Prioritization results obtained from a partially masked eBird data set, restricted to only the species and geographical domain covered by Frei et. al. The eBird results include data from all priority seasons, as identified in the primary manuscript. The Frei et. al. products only cover breeding populations, so no other seasonal distributions are included in those maps.

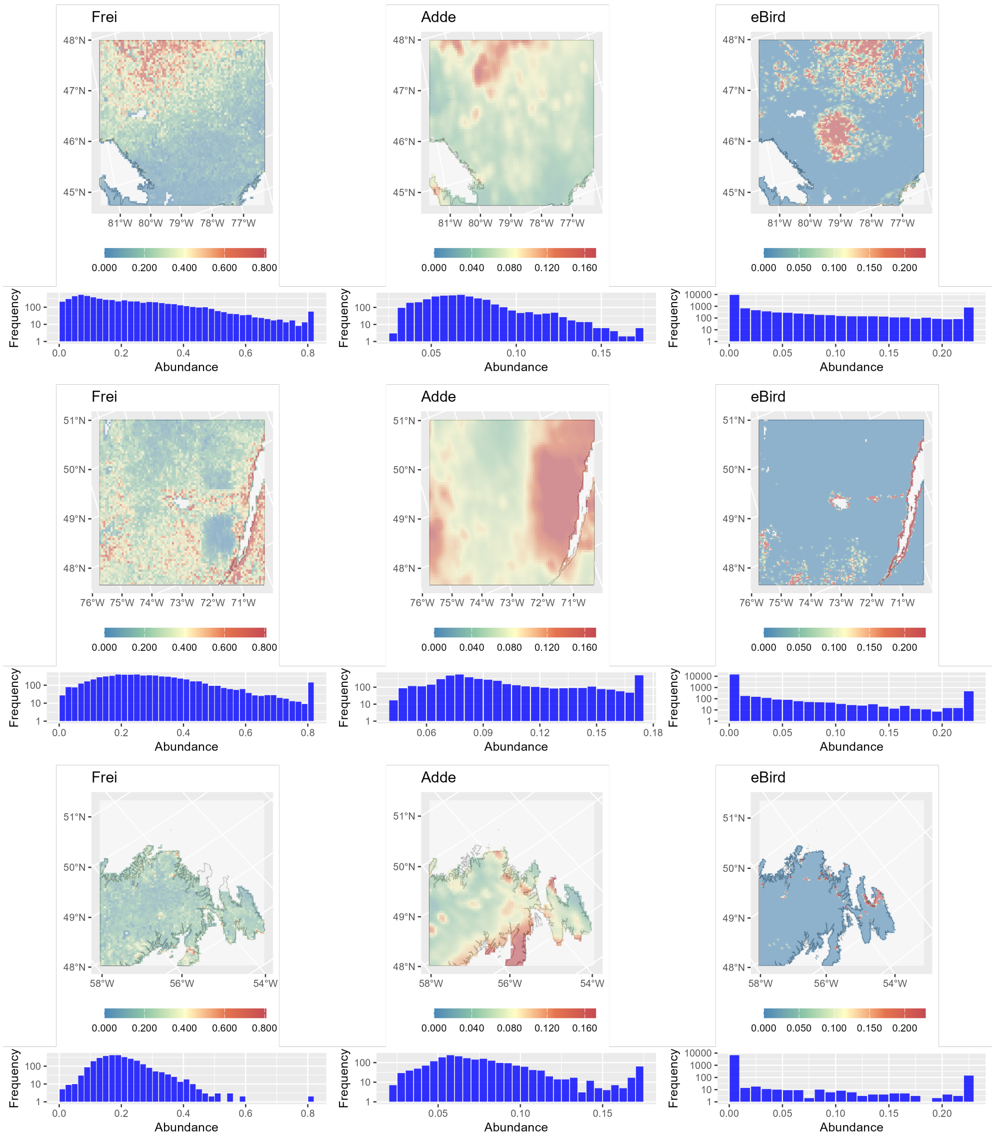


**Fig. E14**: Comparison of three sources of abundance data ((Frei et al. 2024; Adde et al. 2020; Fink et al. 2025)) for the American Black Duck in different regions of the EHJV. Top: Central Ontario, centered on Algonquin Provincial Park. Middle: Central Quebec, including the St. Lawrence estuary. Bottom: The eastern coast of Newfoundland

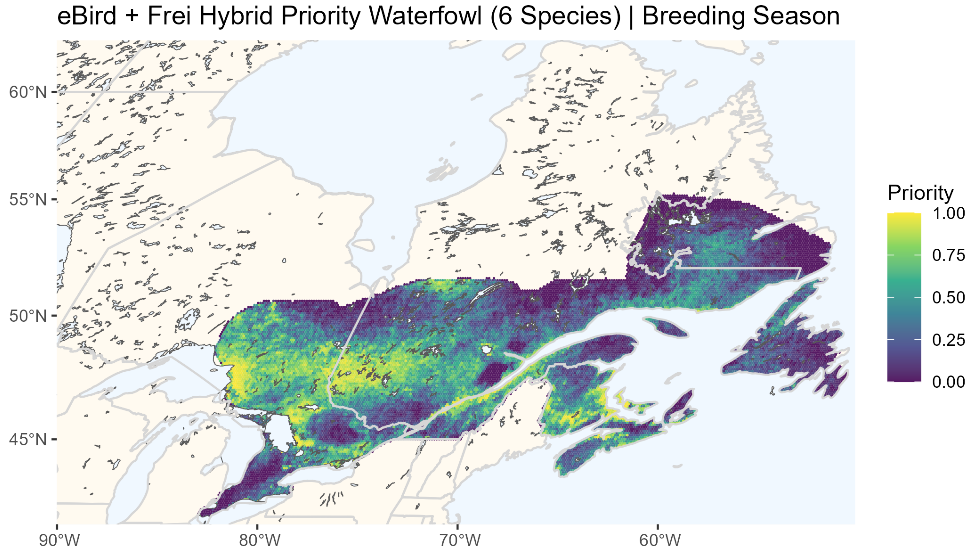


**Fig. E15**: Prioritization results based on a hybrid collection of abundance layers from Frei et. al. (for American Black Duck, Common Goldeneye, Mallard, and Wood Duck in their respective breeding seasons) and eBird Status and Trends (for the other priority seasons of these four species, as well as all priority seasons for the Barrow's Goldeneye and Common Eider).

### Appendix F Potentially Restricted Areas

As a reference, the EHJV-wide prioritization maps are reproduced here with overlays indicating lands which may not be available for conservation delivery, depending on the organizations and delivery mechanisms in question. Grid cells are occluded if their area is covered by at least 50% of either of two classes:

1. Lands held in reserve by the Federal government for military or other purposes, based on data from the Canadian Forest Service's Map of Forest Management in Canada (Government of Canada; Natural Resources Canada 2020).
2. Indigenous lands, as demarcated by Natural Resources Canada's Aboriginal Lands of Canada Legislative Boundaries map, which demarcates "legislative boundaries of Indian Reserves, Land Claim Settlement Lands (lands created under Comprehensive Land Claims Process that do not or will not have Indian Reserve status under the Indian Act) and Indian Lands" (Government of Canada; Natural Resources Canada 2024).

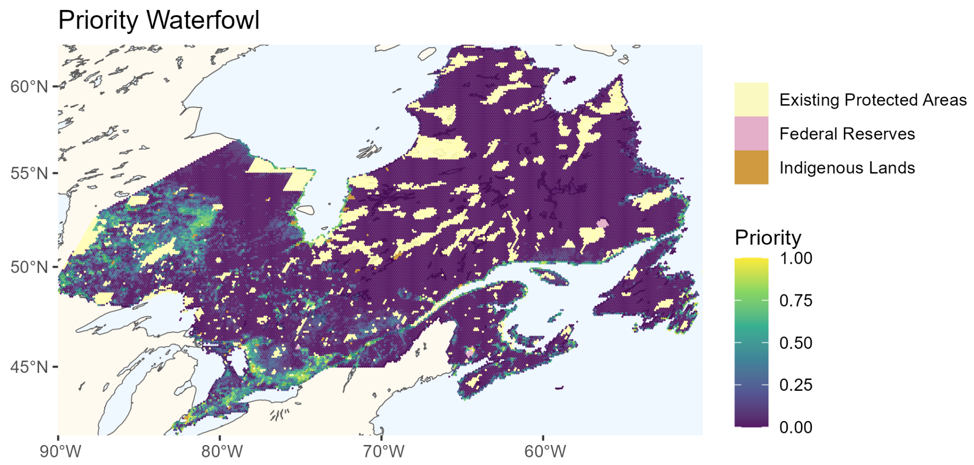


**Fig. F16**: Prioritization of target waterfowl species, overlaid to show grid cells belonging to potentially unavailable classes.

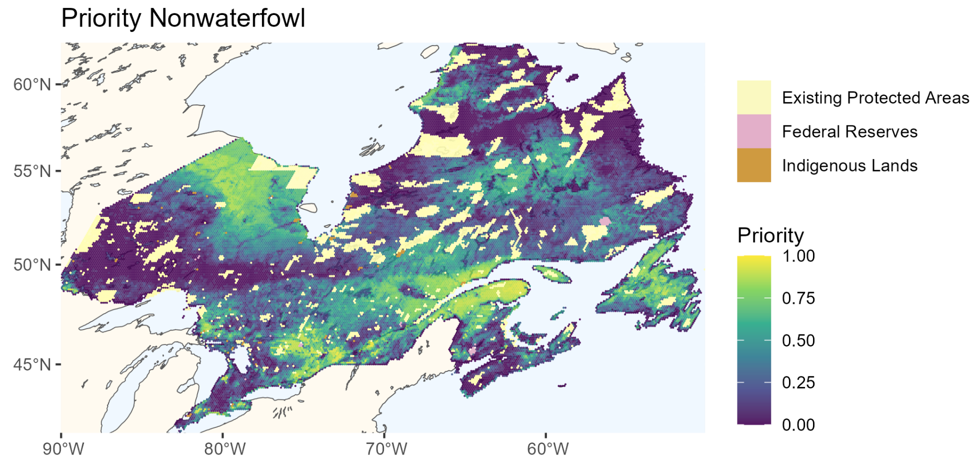


**Fig. F17**: Prioritization of target non-waterfowl species, overlaid to show grid cells belonging to potentially unavailable classes.
